# A unified analysis of atlas single cell data

**DOI:** 10.1101/2022.08.06.503038

**Authors:** Hao Chen, Nam D. Nguyen, Matthew Ruffalo, Ziv Bar-Joseph

## Abstract

Recent efforts to generate atlas-scale single cell data provide opportunities for joint analysis across tissues and across modalities. Most of the existing methods for single cell atlas analysis use cells as the reference unit to combine datasets. However, such methods suffer from the inability to effectively integrate cross-modality data, hindering downstream gene-based analysis, and loss of genuine biological variations. Here we present a new data integration method, GIANT, which is for the first time designed for the atlas-scale analysis from the gene perspective. GIANT first converts datasets from different modalities into gene graphs, and then recursively embeds genes in the graphs into a latent space without additional alignment. Applying GIANT to the HuBMAP datasets creates a unified gene embedding space across multiple human tissues and data modalities, where gene representations reflect the functions of genes in their cells. Further evaluations demonstrate the usefulness of GIANT in discovering diverse gene functions, and underlying gene regulations in cells of different tissues.

## Background

Several recent efforts have attempted to generate atlas-scale single cell data. Examples include The Human BioMolecular Atlas Program (HuBMAP) [1], the Human Cell Atlas (HCA) [2], the Cellular Senescence Network (SenNet) [3], the Human Lung Cell Atlas (HLCA) [4] and many more. In all of these efforts, researchers aim to collect and analyze data from multiple tissues, modalities, conditions and more. While the initial phases of the programs often focused on the coverage of different tissues, and on the separate, modality-specific analysis of each dataset, we are now at a point where, for the first time, we can perform joint cross-tissue and cross-modality analysis, on single cell data at the atlas scale.

The joint analysis is challenging for many reasons. First, one needs to decide on a common reference unit for the analysis. While most of the prior methods for atlas analysis focused on cells [5–8], such analysis often fails to combine data across modalities. In many cases, the large technical variation between data modalities can lead to inappropriate matching cells being selected between datasets, which results in ineffective integration [9]. In addition, cell based analysis makes it more difficult to identify shared pathways between different cell types, in part because it is not focused on the gene space [10]. Another issue is the type of information the method can handle. Cell embedding and clustering based on gene expression similarity, which has been commonly used [11–13], may not be appropriate for spatial technologies which provide information on cell relationships in space [14–16]. Finally, different signal-to-noise levels across datasets [10], and the large volume of data in atlas studies create considerable challenges for joint analysis.

To date, there have been some attempts at a large-scale joint analysis of data from different modalities (not just in biology). The main approach used for such integration utilizes alignment-based multi-view learning [7, 17–19]. The goal is to transform multi-view (multi-dataset) inputs, represented as a multimodal distribution, into a homogeneous unimodal distribution for down-stream learning tasks. The key idea of these methods is to project the manifolds of multiple datasets into a common space such that both, the locality of each dataset is preserved and the distance between the same entities in different datasets (for example, the same individuals in different social networks) are minimized. While this may work for some biological datasets, we often observe over-alignment problems with this approach [10]. Some other methods require paired data (multimodal measurements derived from the same set of samples) to learn the mapping between different modalities [20, 21], which are usually not available in real cases.

To improve on these issues and to enable large-scale joint analyses of atlas-level data, we developed a new method GIANT (Gene-based data Integration and ANalysis Technique), that focuses on genes rather than on cells. Genes provide a level of granularity which on the one hand can serve as anchor points common to all modalities, while on the other hand makes it easy to perform higher-level analysis. We first convert cell clusters from each dataset and each modality into gene graphs based on the specific characteristics of that modality. Genes are more basic building blocks for joint analysis than cells or pathways and, as we show, such graphs can be extracted from different high-throughput methods that are being profiled in large atlas studies. To integrate these graphs across modalities and tissues, we project genes from all the graphs into a latent space (embedding) using recursive projections. In recursive projection [22, 23], a dendrogram is used to enforce similarity constraints across graphs while still allowing genes with multiple functions to be projected to different locations in the embedding space. In other words, the recursive method can be seen as a hierarchical alignment along embedding, in which graphs nearby in the tree structure are embedded closely in the latent space.

We applied the method to the HuBMAP data from 10 different tissues, 3 different major modalities, and 9 different single cell platforms. We also validated it on another a human development cell atlas dataset [24, 25]. As we show, for both datasets, despite the considerable heterogeneity, our joint processing and analysis method was able to successfully group genes based on their functions and pathways, associate specific pathways with tissues, discover new functions for a large number of genes, and identify tissue-specific regulations of genes based on their cross-modality clustering.

## Results

We developed GIANT to construct a unified embedding for genes from 10 different tissues, including Heart, Kidney, Large Intestine, Liver, Lung, Lymph Node, Pancreas, Small Intestine, Spleen, and Thymus, and 3 different data modalities, which include single cell RNA-seq (scRNA-seq), single cell ATAC-seq (scATAC-seq), and spatial transcriptomics (Slide-seq). Figure 1 presents an overview of GIANT. We first construct gene graphs for cell clusters from each tissue and data modality based on expression or epigenetic correlations (Methods). Next, all gene graphs are connected in a cell cluster dendrogram. A recursive graph embedding method which takes into account both the dendrogram and gene identity connections is applied. The final result is an embedding (a location in a latent space) for each gene in each of the graphs. See “Methods” section for details.

**Figure 1:**
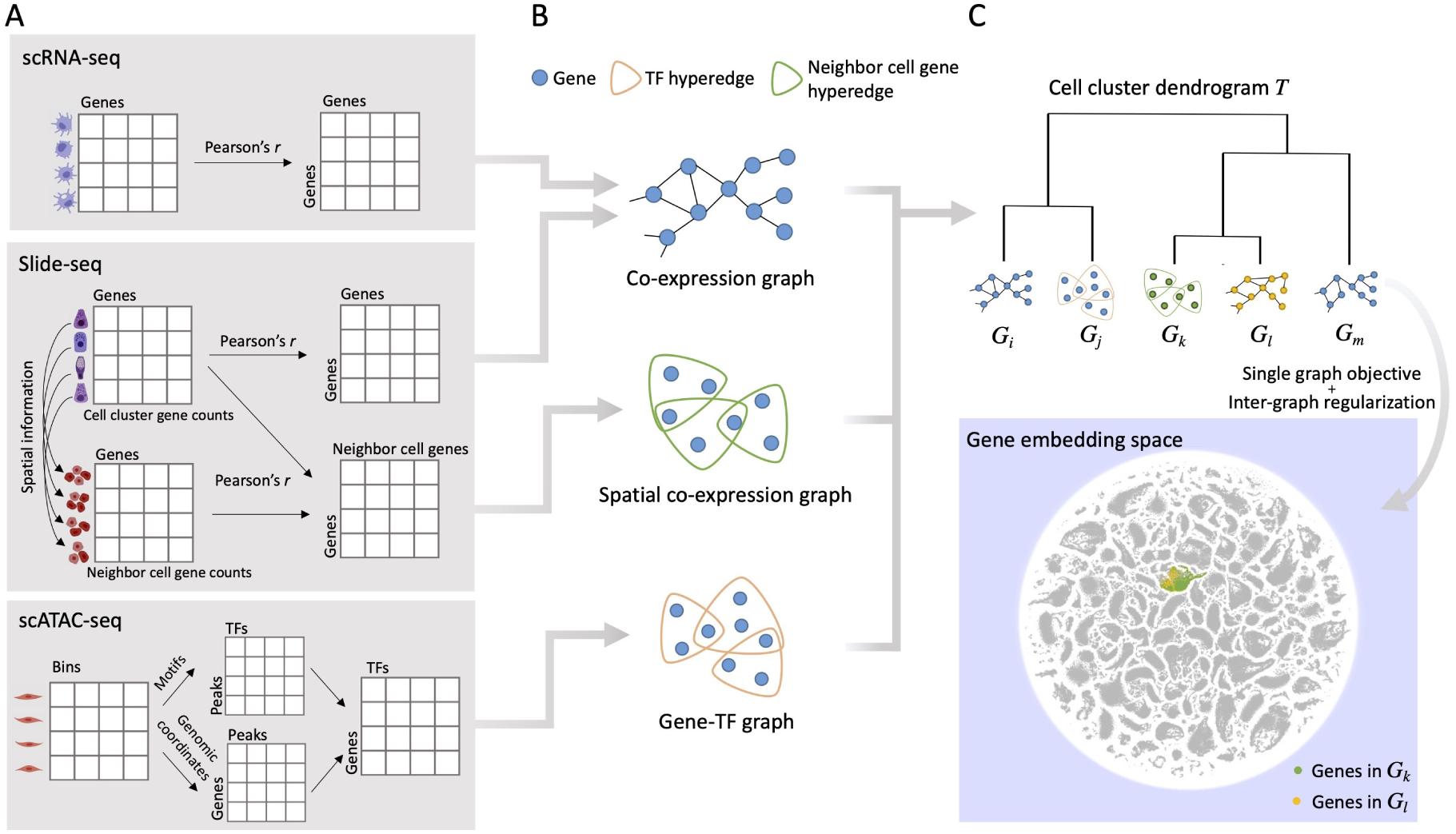
Schematic illustration of the method. **A** Cell clusters are identified from each tissue of each data modality. For each scRNA-seq cell cluster, a gene co-expression matrix is constructed for all the gene pairs in the cell cluster. For each Slide-seq cell cluster, the same gene co-expression matrix is also constructed, and another spatial co-expression matrix is built for gene pairs between the cells of this cell cluster and their neighboring cells. For each scATAC-seq cell cluster, identified peaks from the bin matrix are associated with TFs and genes, which forms a gene-TF matrix. **B** We obtain gene co-expression or co-regulation graphs for each of these modalities based on the specific matrices computed in **A**. **C** A dendrogram for graphs in **B** is constructed by hierarchical clustering of cell clusters, and a multi-layer graph learning algorithm is applied using the tree structure. Each gene in each graph is then encoded into an embedding space, where genes with similar functions are embedded together.

### GIANT integrates multi-modality and multi-tissue data

The gene embedding space produced by GIANT enables joint functional analyses across data modalities, tissues, and cell clusters. Therefore, the location of a gene in the space is expected to reflect the function the gene performs (which could be the same across several different cell types and tissues), instead of the specific data source which generates the value for that gene. For example, genes from different tissues can be closely embedded in space as long as they perform similar functions in these tissues. To assess the integration of various data sources, we visualized the entire embedding space and highlighted the locations of genes based on their respective data sources. Figure 2 presents the distribution of genes in the map based on their modalities (Fig. 2A), tissues (Fig. 2B), and cell clusters (Fig. 2C) respectively. As can be seen, the locations of genes are not determined by their data modalities or tissues, allowing us to use the embeddings to investigate shared functionalities across tissues and mechanisms being studied by each of the data modalities we used. This observation is confirmed by our quantitative evaluation. GIANT obtains a very low silhouette coefficient (0.016) for data modalities (Supplementary Notes). When compared to six cell-based data integration methods that are suitable for the data we used (Supplementary Notes, Supplementary Figures 1-2), GIANT achieves a substantially lower silhouette coefficient of data modality than Harmony [26], LIGER [27], Scanorama [18], scVI [28], and GLUE [29], and a similar silhouette coefficient to Seurat [6] (Supplementary Figure 1G), where LIGER, GLUE, and Seurat are specifically designed for integrating data across multiple modalities. In addition, for the scRNA-seq data used, which were obtained from five different platforms, we see very good integration by GIANT (Supplementary Figure 3).

**Figure 2:**
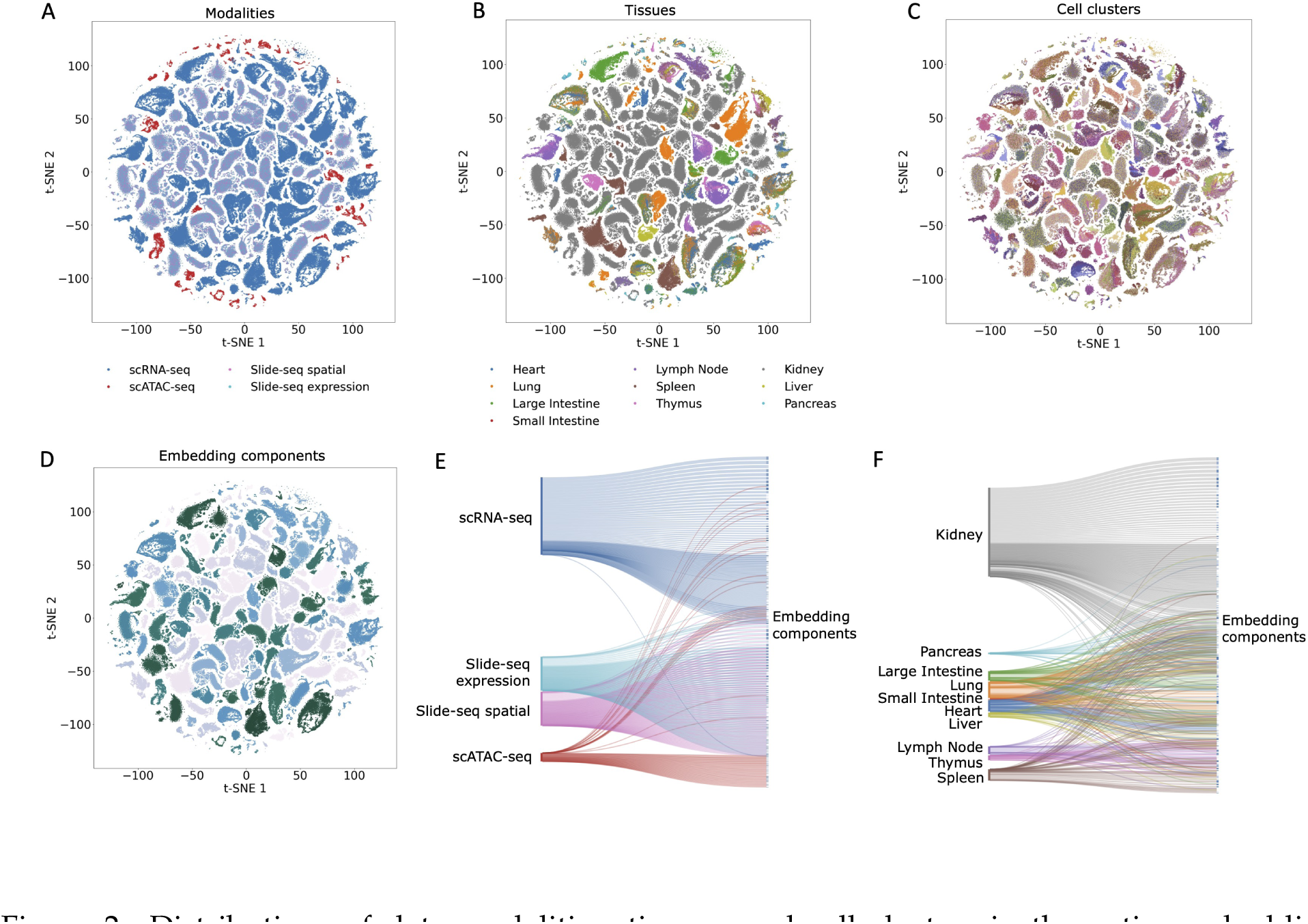
Distributions of data modalities, tissues, and cell clusters in the entire embedding space. **A** Distributions of data modalities in the embedding space, where genes are colored by data modalities. **B** Distributions of tissues in the embedding space, where genes are colored by tissues. **C** Distributions of cell clusters in the embedding space, where genes are colored by cell clusters. **D** The embedding space is split into components, where genes in different components are shown in different colors. **E** Cross clustering between data modalities. **F** Cross clustering between tissues. As can be seen, genes from several different tissues, and from multiple modalities, are clustered together, allowing for a joint analysis of this large dataset.

To further investigate which data sources tend to co-cluster, we clustered the entire embedding space (Fig. 2D), and then looked at the data sources assigned to each of the resulting clusters (embedding components). Specifically, we applied the Leiden algorithm [30] to cluster all genes in the embedded space into 172 components (Methods). We then determined the data modality or tissue for each gene in each component. Figure 2E shows the mapping from data modalities to components. Genes from Slide-seq expression and Slide-seq spatial graphs are always assigned to the same components, as these represent different measurements for the same gene. For the other modalities, we found that 14.5% (25 out of 172) of the components contain genes from at least two of the three modalities, scRNA-seq, scATAC-seq, and Slide-seq. Figure 2F shows the mapping from tissues to components. Interestingly, genes from Heart (blue) and Lung (orange), both closely related to the circulatory and the respiratory systems [31], frequently appear in the same components. Tissues from the immune system, including Lymph Node (purple), Thymus (pink), and Spleen (brown) [32], are also frequently co-clustering. These results combined show that GIANT successfully integrated data from different modalities, tissues, and cell clusters, and is able to group genes based on common functions rather than data sources.

### Genes closely embedded in the space show tissue-specific GO enrichment

We next tested the functional enrichment of nearby genes in the embedding space (Methods). Due to the large number cells of different tissues we analyzed, such analysis allowed us to identify functional groups spanning a large portion of the Gene Ontology categories in the embedding components. Specifically, this analysis identified 272,848 enriched Biological Process (BP) GO terms, 44,952 enriched Cellular Component (CC) GO terms, and 56,199 enriched Molecular Function (MF) GO terms (adjusted *P ≤* 0.01) in the 1,199 embedding components (Supplementary Table 1).

We next analyzed the GO terms discovered in the embedding components associated with different tissues. Specifically, for each component, we looked at the top three tissues for genes assigned to that component. Next, we selected the top 5% enriched BP GO terms with the largest fold changes and adjusted *P ≤* 1e *−* 5, and associated these GO terms with each of the top three tissues. To summarize the enriched GO terms for each tissue, we propagated these GO terms to higher levels along the GO hierarchies. The top GO categories enriched in each tissue are shown in Figure 3A, where the size of each dot indicates the ratio of child terms of the noted GO category in all the enriched terms of the tissue, while the color of each dot indicates the fold change of the ratio over the average value of the ratios across tissues.

**Figure 3:**
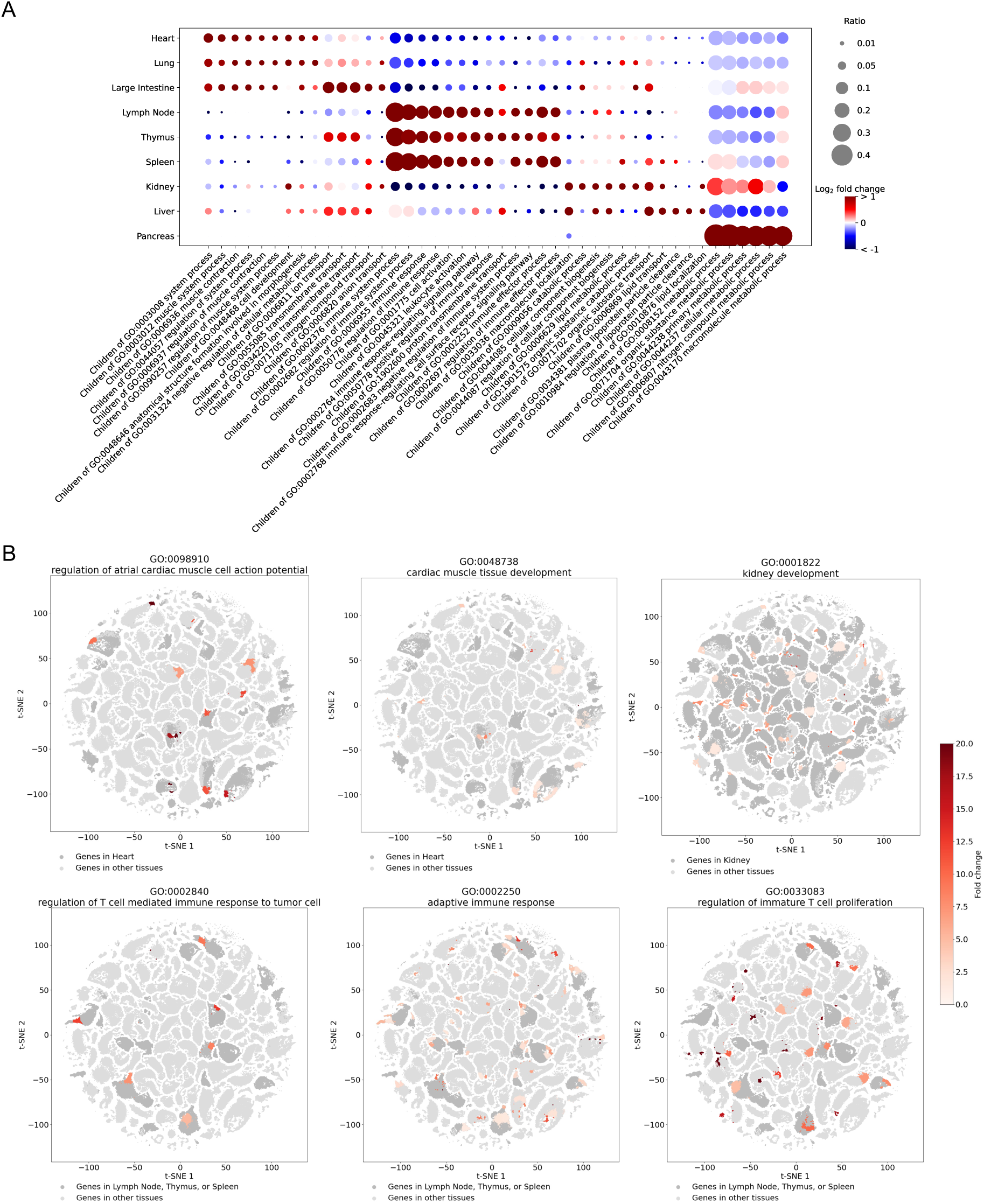
GO terms enriched in different embedding components. **A** The top categories of GO terms enriched in the embedding components associated with different tissues. The size of each dot indicates the ratio of child terms of the GO category in all the enriched terms of the tissue. The color of each dot indicates the fold change of the ratio over the average value of ratios in the column. **B** GO enrichment in different embedding components. Components with the GO term enriched (adjusted *P <* 0.01) are colored by the enrichment levels of the GO term, while the other components are shown in grey. Genes from tissues related to the GO term are highlighted by dark grey, while genes in the other tissues are colored by light grey.

We found that most of the top GO categories for each tissue (dark red dots in Fig. 3A) are strongly consistent with the known cellular functions of that tissue. For example, muscle related GO categories (*e.g.*, GO:0003012 muscle system process, GO:0006936 muscle contraction) are prominent for Heart, Lung, and Large Intestine, which contain abundant cells of cardiac muscles, respiratory muscles, and smooth muscles [33–35]. For immune tissues (Lymph Node, Thymus, and Spleen), immunity related GO categories dominated the assignments, including GO: 0002376 immune system process, and GO:0006955 immune response. Lipid metabolism related GO categories (*e.g.*, GO:0010876 lipid localization) are prominently discovered for Liver, and metabolic process related GO categories are significantly found for Pancreas, all consistent with the specific functions of these tissues [32, 36, 37]. Examples of some tissue-specific GO terms are shown in Figure 3B. Each of them is enriched in different components in the embedding space (reddish regions), while they are more likely found in components where the related tissues reside in (dark grey regions).

We again compared GIANT with the six aforementioned cell-based embedding methods. WGCNA module detection method [38] was applied in each cell cluster identified from each cell-based embedding (Supplementary Notes). We found that compared with GIANT, there are much fewer unique GO terms enriched in the gene modules discovered from cell-based embeddings (Supplementary Figure 1G), suggesting that GIANT performs better in discovering functional gene modules across datasets.

### Testing GIANT integration on a developmental atlas dataset

To further test the ability of GIANT to integrate data from different modalities, we used a human fetal cell atlas dataset [24, 25]. This dataset contains single cell data from 15 tissues and 2 data modalities, including Adrenal, Cerebellum, Cerebrum, Eye, Heart, Intestine, Kidney, Liver, Lung, Muscle, Pancreas, Placenta, Spleen, Stomach, and Thymus of scRNA-seq and scATAC-seq. Unlike the HuBMAP data, the fetal atlas provides cell type annotations for cell clusters, allowing us to validate our embeddings at a more refined level. Results are presented in Figures 4A-B (Supplementary Notes). As the figure shows, GIANT is able to merge genes from different modalities and tissues as seen for the HuBMAP dataset. Specifically, several components in the embedding space are dominated by one cell type (Fig. 4C), though gene in these components are often from several tissues and both modalities.

**Figure 4:**
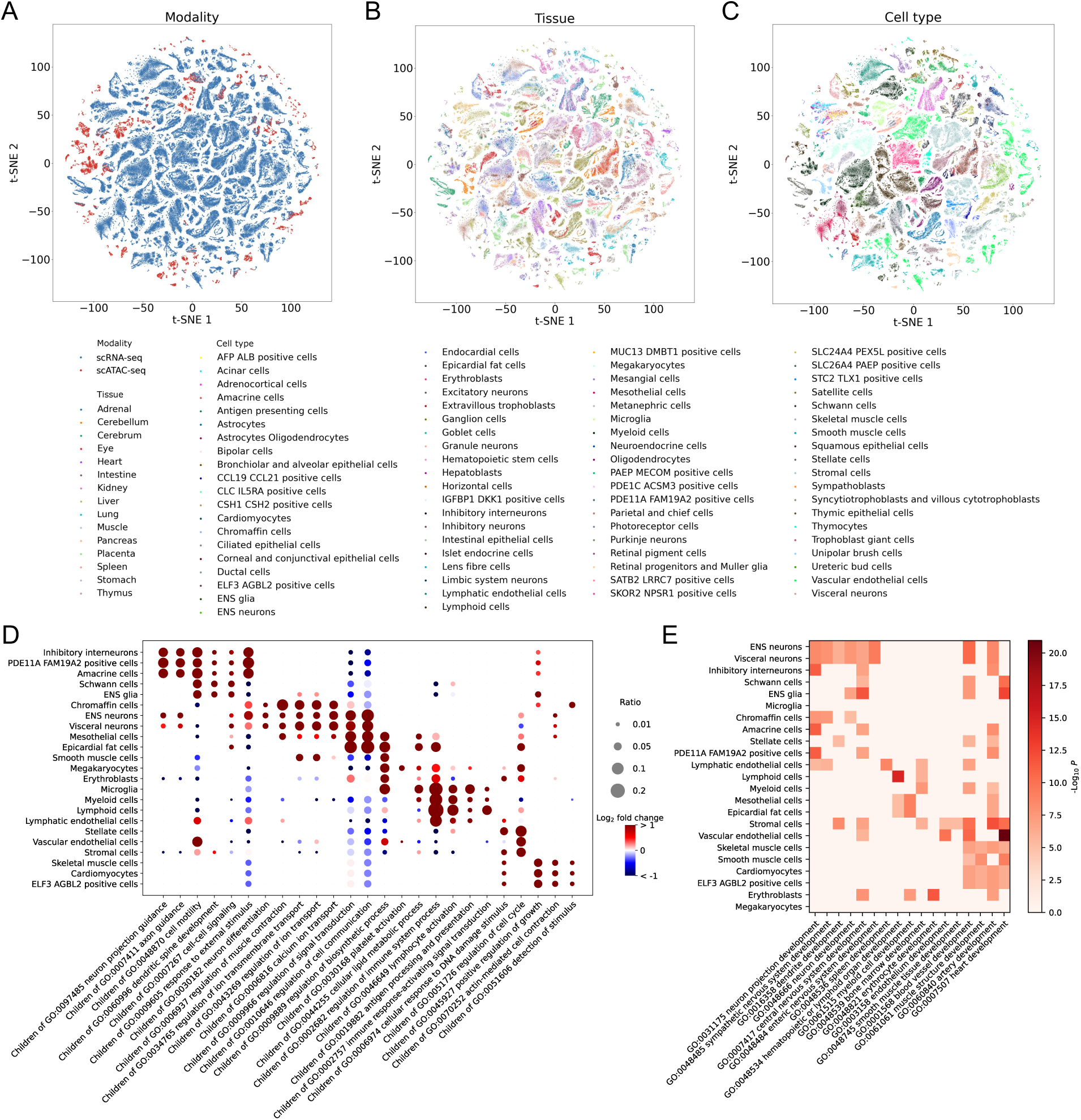
Application of GIANT on the human fetal atlas. **A** Distributions of data modalities in the embedding space. **B** Distribution of tissues in the embedding space. **C** Distribution of cell types in the embedding space. Each point in the plots represents a gene that is colored by its data source. **D** Top GO categories for different tissues. The size of each dot indicates the ratio of child terms of the GO category in all the enriched terms of the tissue. The color of each dot indicates the fold change of the ratio over the average value of ratios in the column. Shown cell types are those with at least 50 GO terms. **E** The enrichment level of different developmental GO terms in the embedding components associated with different cell types shown in **D**. Colors show the negative logarithm of the corrected *P* values.

We also performed GO enrichment analysis in the embedding components, this time using cell types instead of tissues. Our analysis revealed a strong consistency between the enriched Gene Ontology (GO) terms found within the embedding components of each cell type (Fig. 4D) and the well-established cellular functions associated with those specific cell types. For example, signal transduction related GO categories (*e.g.*, GO:0007267 cell-cell signaling, GO:0009966 regulation of signal transduction) are mainly assigned to nerve cells, such as inhibitory interneurons [39], and enteric nervous system (ENS) neurons [40], while immune related GO categories (*e.g.*, GO:0002682 regulation of immune system process, GO:0046649 lymphocyte activation) are mostly associated with immune cells, such as myeloid cells [41], lymphoid cells [42], and microglia [43]. We also found many development related GO terms enriched in embedding components for this fetal atlas dataset. A closer examination revealed that the cell type-specific developmental pathways are highly enriched in the embedding components associated with the corresponding cell types (Fig. 4E), such as neuron projection development (GO:0031175) for inhibitory interneurons, hematopoietic or lymphoid organ development (GO:0048534) for lymphoid cells, and erythrocyte development (GO:0048821) for erythroblasts.

Finally, we found that 88.5% (1,061 out of 1,199) of the HuBMAP embedding gene components are enriched in the embedding components of the human fetal dataset (Supplementary Figure 4), indicating the ability of large scale atlas datasets to reveal most of the significant biological processes.

### Inferring gene functions from multi-modality multi-tissue embeddings

The precise function of a specific protein (and the genes that code for it) may differ between tissues and cell types [44]. Given the observations above that GIANT can group functionally similar genes together across data sources, we used the embeddings to infer novel gene functions for all the genes we embedded. We first clustered each gene’s embeddings and then inferred the gene’s functions in each cluster based on other co-clustered genes with known functions in the graphs of this cluster (Methods).

We tested the coverage and precision of the functions that we can assign using GO. We found that 47.4% of all known GO functions for the genes are identified in at least one of their clusters, which means that our predictions cover roughly half of all known functions for these genes (recall). This number is much higher than the expected number given the number of GO assignments (compared to 6.9% expected for random assignments of 671,215 GO terms for the 1,822 genes analyzed here). On the other hand, the ratio of known functions in our GO assignments for genes (precision) is 25.1% which is also much higher compared to the ratio of known functions in the whole prediction space (1.5%) (Fig. 5A). Precision and recall of our predictions considering only child terms of other GO categories are shown in Figure 5A.

**Figure 5:**
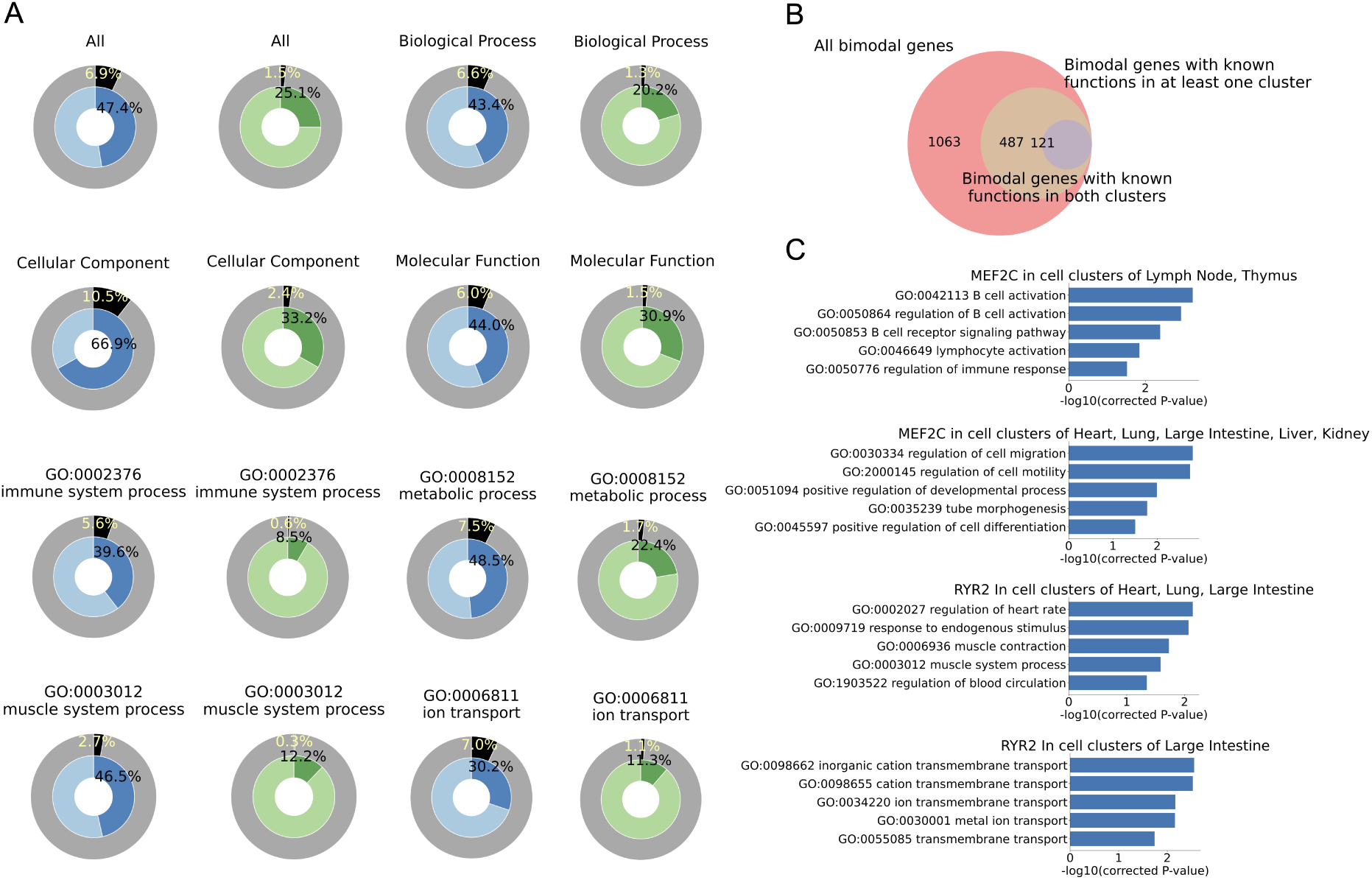
Inference of gene functions in cell clusters. **A** Blue circles show the ratios of known gene functions that are covered by our predictions (recall), while their outside circles indicate the ratios of our predictions in the whole prediction space (all the possible GO assignments). Green circles show the ratios of known gene functions in our predictions (precision), while their outside circles indicate the ratios of known functions in the whole prediction space. Titles indicate child GO terms of which categories are considered. Besides the whole GO space and three main GO branches, some representative GO categories in Figure 3A are also shown here. **B** The number of bimodal genes, and the number of bimodal genes for which one or both functions are confirmed. **C** Inferred functions of two genes in different cell clusters.

Given this large coverage and precision, we next investigated genes that are assigned very different functions in at least two different clusters, which are termed as “bimodal genes” here. In particular, genes are selected as bimodal that met the following criteria: among all the gene’s clusters with at least 5 assigned GO terms, which are BP terms associated with no more than 200 genes to focus on specific GO terms that may show tissue specificity, at least two clusters do not have any common GO terms assigned. If a gene has more than two such clusters, the two with the most GO terms assigned are considered in this analysis. As a result, we identified 1063 bimodal genes, and 487 (45.8%) among them have known functions assigned in at least one cluster. For 121 (11.4%) of them, both clusters have functions that are already known even though they can be very different (Fig. 5B).

Figure 5C presents examples of bimodal genes with different known functions in different clusters. MEF2C is essential in B-lymphopoiesis, as well as for B-cell survival and proliferation in response to stimulation [45, 46]. On the other hand, it is known to be involved in vascular development and controls cardiac morphogenesis [47, 48]. Our method correctly assigns MEF2C the functions of B cell activation, lymphocyte activation, and regulation of immune response in the group of Lymph Node and Thymus cell clusters, while it assigns this gene the functions of positive regulation of developmental process, and tube morphogenesis in a group of Heart and Lung cell clusters. Similarly, RYR2 has been found to play an important role in triggering cardiac muscle contraction and to be related to cardiac arrhythmia [49], while it also mediates the flow of calcium ions across cell membranes in large intestine [50]. Again, RYR2 is assigned the functions of regulation of heart rate, muscle contraction, and regulation of blood circulation in the group containing a Heart cell cluster, while in the Large Intestine cell cluster, it is assigned to the ion transmembrane transport function. A complete list of predicted bimodal genes and their predicted functions can be found in Supplementary Table 2.

### Gene embeddings enable the discovery of tissue-specific gene regulations

Identifying the regulatory programs of cells helps understand their diverse gene expression patterns. We further demonstrate the ability of our gene embeddings to discover gene regulations in cell clusters of different tissues. Similar to the previous subsection, we calculated the enrichment of genes regulated by the same transcription factor (*i.e.*, TF regulon) in each embedding component (Methods). Specifically, this analysis identified 5,815 enriched TF regulons (adjusted *P ≤* 0.001) in the 1,199 components. To compare the TFs identified in cells of different tissues, we associated discovered TFs with tissues using the top three tissues for each component as discussed above. TF overlaps among different subsets of tissues are shown in Figure 6A. Although a large number of TFs (104 out of 435, 23.9%) were identified as significant for all the tissues, most TFs show specificity for different tissues. These observations are consistent with previous findings in the literature [51, 52]. The full list of TFs identified in different tissues can be found in Supplementary Table 3.

**Figure 6:**
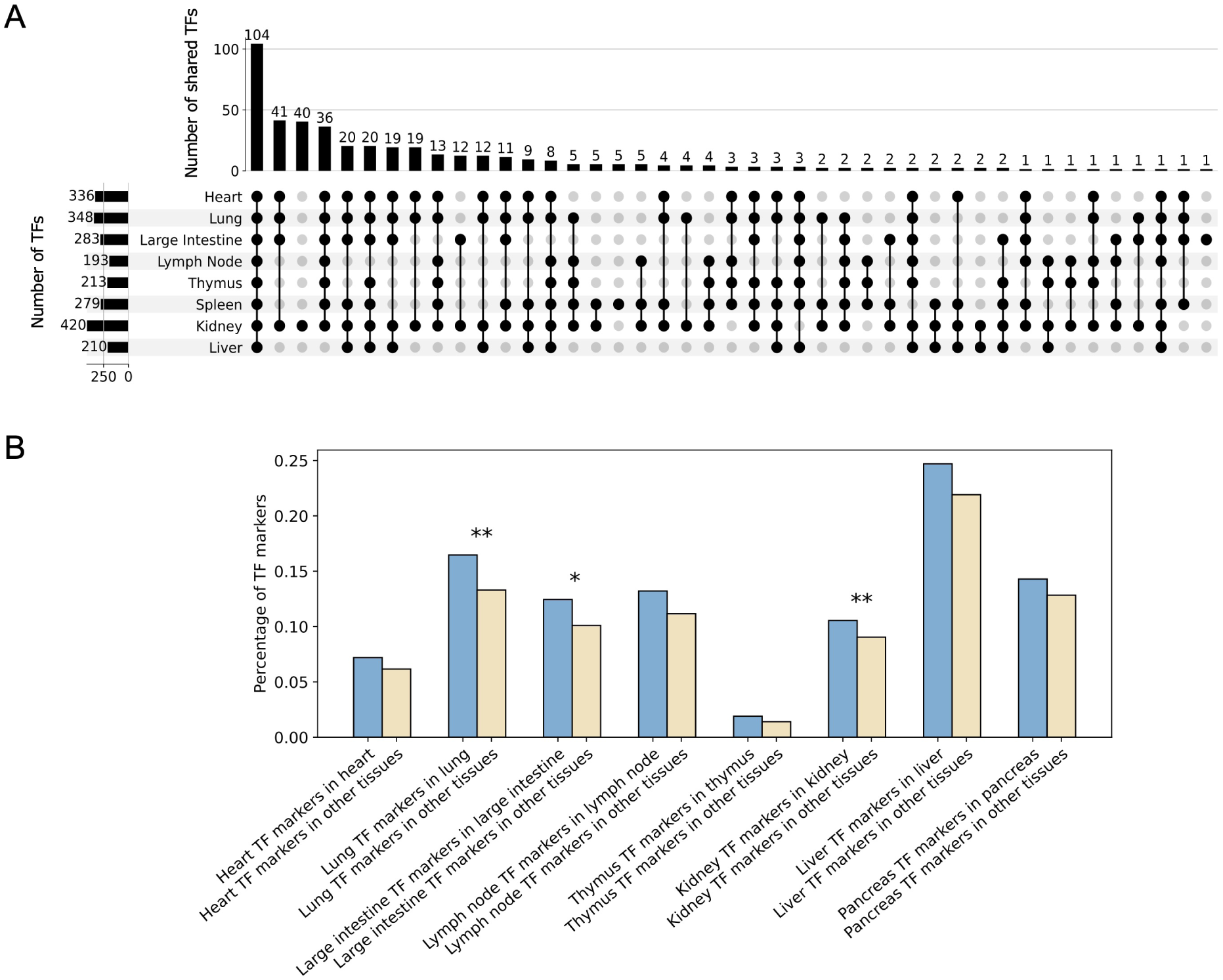
TFs discovered in different tissues. **A** TF intersections among different tissues. Bars on the left show the number of TFs identified in each tissue, while each bar on the top shows the number of TFs shared among tissues marked by the dots below it. Tissues with fewer than 20 TFs identified are omitted in this figure. The figure was drawn using UpSetR [70]. **B** The percentage of tissue-specific TF markers in the corresponding tissue (blue) compared to their percentage in the other tissues (light yellow). Significant enrichment in blue groups are marked by stars (*: *P ≤* 0.05, **: *P ≤* 0.01, *P*-values are evaluated using one-sided Fisher’s exact tests).

To further validate our discoveries, we examine whether TFs found in each tissue are enriched with TFs known to be specific to this tissue. For this, we used a prior list of tissue marker TFs [53]. We find enrichment for tissue markers in all tissues and significant enrichment for three of the eight tissues (Fig. 6B).

## Discussion

Recent efforts to generate atlas-scale single cell data provide opportunities for joint analysis across tissues and modalities, which can enable new insights into cell functional plasticity, gene function, cell regulation, and disease. A key challenge for such analysis is how to effectively integrate data from different sources, such as tissues, modalities, and conditions, while enabling the discovery of system-level organizations and regulations.

We developed a new method which is the first to fully integrate atlas-level data from multiple tissues and modalities in a single unified embedding. For this, we use genes as the anchor points common to all datasets, modalities and tissues. Our method first converts cell clusters in different tissues from different data modalities to gene graphs and then recursively embeds these graphs such that connected genes within a single graph are encouraged to have similar embeddings, while genes from graphs generated from similar cells are projected closely in the embedding space. Finally, a unified embedding space is produced where the locations of genes reflect the functions they perform in the cells.

We applied our method to HuBMAP database with 10 different tissues, 3 different major modalities and 9 different single cell platforms. As we have shown, despite the considerable heterogeneity of the input data, our method was able to successfully group genes based on their function and pathways. We further used the method to assign new functions to previously characterized genes, and explore tissue similarities and regulations.

An interesting finding is the number of genes that have at least two sets of functions (termed “bimodal genes”). Even with only 10 tissues, we identified over 1000 genes with two or more diverse functions depending on the tissues. While several studies have pointed out genes with multiple functions [54] mostly in different cell types or tissues [45, 46], these prior findings were anecdotal and focused on very few genes. By analyzing much larger atlas data we were able to determine that such bimodal genes are actually quite common. It is likely that the number of genes with multiple functions is actually much larger given the large number of tissues that were not included in our study. These findings indicate that even though the majority of genes have at least one associated function in large annotation databases (for example, GO), there is still much work needed to fully characterize the functions of human genes.

For HuBMAP we have mainly focused on tissue-level analysis as the first step for atlas-level integration. While we used cell clustering (which likely represents cell types) as input, these clusters were not assigned to specific cell types and so our method can currently only provide information on tissue-level functions and regulations. However, as we showed with the fetal atlas dataset, GIANT can be easily applied to integrate atlas data at the cell type level as well.

While GIANT relies on cell-based clustering for the initial processing (creating dataset specific graphs), it integrates data in the gene space rather than at the cell level. This provides several advantages as we discussed above and leads to better results as we showed.

GIANT constructs gene-TF graphs for scATAC-seq cell clusters, which connect genes to TFs who are known to bind to their promoter regions while the binding sites are open. This reduces the scATAC-Seq data to the gene space which, on the one hand, leads to information loss but on the other enables easier integration of data from different modalities.

GIANT successfully integrates data from three different modalities (the silhouette coefficient for data modality is 0.016 indicating very little impact of data modalities, Supplementary Figure 1G). However, gene nodes from individual modality graphs (scATAC-Seq, scRNA-Seq *etc.*) are usually not merged since they are embedded together as part of their input graph. This is likely due to the fact that graphs for different modalities have very different structures that capture gene functions from different perspectives (co-expression for scRNA-seq, spatial co-expression for Slide-seq, while co-regulation for scATAC-seq). Still, the ability to integrate gene modules (or graphs in our representation) of GIANT is important and, as we show, can lead to relevant and novel biological findings. For example, for the gene function prediction task, among the 664 genes that appear in at least two of the three data modalities, 332 (50.0%) genes have predicted functions arising out of neighbors from more than one modality. Among the 1,924 genes that appear in at least two tissues, 1,606 (83.5%) genes have predicted functions arising out of neighbors from more than one tissue (Supplementary Figure 5). We also note that GIANT can enforce a higher level of modality integration by increasing the regulation strength (*λ*), however, in such cases we observed lower-quality gene embeddings (Supplementary Figure 6). Similarly, we found that cell-based embeddings, such as Seurat and GLUE, achieve higher levels of modality integration, but the resulting gene modules are of lower quality (Supplementary Figure 1).

There are other methods that can also generate gene embeddings from single cell data, such as GLUE [29], DeepMAPS [55], and SIMBA [56], however, such methods only generate one embedding for each gene in the whole dataset. In contrast, GIANT generates one embedding for each gene in each of the cell clusters, which reflects the functional variability of the genes in tissues or cell types.

### Conclusions

Our embedding method is implemented in Python and is available along with the generated gene embeddings at https://github.com/chenhcs/GIANT. We hope that as the analysis of atlas-level data becomes more common, our method would be able to be applied to additional atlases and datasets to integrate large-scale single cell data.

## Methods

### Datasets

We integrated single cell data from more than 60 individuals profiling 10 different tissues and 3 major data modalities from the Human BioMolecular Atlas Program (HuBMAP) consortium [1]. **Single cell RNA-seq (scRNA-seq) datasets**: scRNA-seq datasets were obtained for 8 tissues: Heart, Kidney, Large Intestine, Liver, Lung, Lymph Node, Spleen, and Thymus. For Heart we obtained 14 datasets and a total of 30,241 cells, 15 datasets and 109,055 cells for Kidney, 6 datasets and 13,128 cells for Large Intestine, 5 datasets and 7,916 cells for Liver, 9 datasets and 68,390 cells for Lung, 4 datasets and 40,449 cells for Lymph Node, 10 datasets and 93,737 cells for Spleen, 3 datasets and 10,838 cells for Thymus. This data was obtained from 5 platforms (Supplementary Table 4) and uniformly processed. The uniform processing used the Salmon’s Alevin method [57] for sequencing reads mapping and abundance quantification, leading to a cell-by-gene count matrix for each dataset.

#### Single cell ATAC-seq (scATAC-seq) datasets

scATAC-seq datasets were obtained for 8 tissues, including Heart, Kidney, Large Intestine, Liver, Lung, Pancreas, Small Intestine, and Spleen. There are 14 datasets and in total 312,579 cells for Heart, 19 datasets and 272,526 cells for Kidney, 6 datasets and 8,368 cells for Large Intestine, 2 datasets and 50,748 cells for Liver, 9 datasets and 205,048 cells for Lung, 3 datasets and 68,294 cells for Pancreas, 3 datasets and 7,774 cells for Small Intestine, 4 datasets and 16,260 cells for Spleen. Again, data from 3 platforms (Supplementary Table 4) was combined using a uniform processing pipeline. For this, we used SnapTools [58] for reads mapping and abundance quantification. SnapTools segments the genome into uniform-sized bins of 5,000 base pairs (bp) and a cell-by-bin matrix is created for each dataset with each element indicating the count of sequencing fragments overlapping with a given bin in a certain cell.

#### Slide-seq datasets

Slide-seq [59], a spatial transcriptomics method, was used to profile Kidney samples. There are in total 551,342 cells (DNA-barcoded beads) included in 19 Slide-seq datasets. Slide-seq generates short read sequencing data that is equivalent to scRNA-seq data, and was processed in the same way we processed all scRNA-Seq data. In addition, the spatial location associated with each bead is provided.

The HuBMAP data from each modality (even from different platforms) were processed using the same pipeline and the reads of different datasets were aligned to the same reference genome. It is highly recommended that other large-scale atlas analyses follow a similar procedure.

### Clustering cells with the HuBMAP modality-specific pipelines

For each tissue in each data modality, cells are merged by joining the data matrices of different datasets along the cell axis. The HuBMAP modality-specific data analysis pipelines, described below, are then used to cluster cells in each tissue of each data modality.

The HuBMAP scRNA-seq pipeline uses the Scanpy [60] package to perform cell clustering. Scanpy first filters cells that express less than 200 genes and genes that are expressed in less than three cells. Each cell is then normalized by its total counts over all genes. The produced count matrix is log transformed and scaled so that each gene has unit variance and zero mean, after which it is used as input to dimensionality reduction via PCA, computation of a k-nearest-neighbor (k-NN) graph in PCA space, and finally clustering with the Leiden algorithm [30]. We further identify differentially expressed genes for each cell cluster against all the other clusters in the tissue using the Wilcoxon rank-sum test, allowing for estimating similarities between cell clusters.

The scATAC-seq processing pipeline uses the SnapATAC package [58] for data processing, which starts from each cell-by-bin count matrix. High-quality cells are identified for subsequent analysis based on three criteria: (1) total number of unique fragment count [*>*2000]; (2) UMI count [*>*1000]; and (3) mitochondrial read ratio [*<*50%]. Bins are then filtered out if they overlap with the ENCODE blacklist (downloaded from http://mitra.stanford.edu/kundaje/akundaje/release/blacklists/), are in mitochondrial chromosomes, or belong to the top 5% invariant bins ranked by the coverage. From the filtered cell-by-bin matrix, we create a gene activity matrix. Gene coordinates are extracted and extended to include the 2,000 bp upstream region [6]. Gene activities are then computed based on the number of fragments that map to each of these regions, using the “calGmatFromMat” function in the SnapATAC package, and saved as a cell-by-gene matrix. Cell clusters are identified from the cell-by-gene matrix by using it as the input of the same scRNA-seq cell clustering workflow with the steps of normalization and rescaling, dimensionality reduction, k-NN graph computation, and clustering with the Leiden algorithm. After cell clustering, MACS2 [61] is used to call accessibility peaks over all cells for each cluster with at least 100 cells. Genes that are differentially active for each cell cluster against all the other clusters in the tissue are again calculated using the Wilcoxon rank-sum test from the gene activity matrix.

Since Slide-seq yields gene count matrices that are equivalent to those of scRNA-seq, the same cell clustering pipeline that was used for scRNA-seq is applied to Slide-seq data. The spatial information of cells is used to construct the gene spatial graphs as discussed below.

### Constructing gene graphs for cell clusters

Gene graphs are constructed for cell clusters in all tissues and modalities. To focus on cell type specific genes, we filter genes based on their variance in the datasets. Specifically, the union of the 500 most variable genes from each tissue are selected, using the Scanpy package with the criteria defined in Seurat [62]. The variable genes are identified from gene expression matrices for scRNA-seq and Slide-seq, and gene activity matrices for scATAC-seq. The union of these sets results in 7,000 genes that are selected and used for building gene graphs and further analyses.

An unweighted undirected k-NN gene graph is computed for each scRNA-seq cell cluster from its normalized and log transformed expression matrix. For each pair of genes, the Pearson correlation of their expression values across cells in the datasets is computed. Each gene is then connected to the top 10 other genes with the strongest correlation.

A gene-transcription factor (TF) hypergraph is constructed for each scATAC-seq cell cluster by associating the peaks identified in each cell cluster with TFs and target genes. We identify peaks that contain TF motifs using the motifmatchr package which provides information on 633 Human TF binding motifs collected from the JASPAR 2020 [63] database. A binary peaks-by-TF matrix is generated, where an element “1” indicates that the TF’s motifs are found in the region covered by the corresponding peak. A complimentary gene-by-peak matrix is generated, where elements are set to “1” if the corresponding peaks are within the promoter regions of the corresponding genes. Here the promoter region is defined as the 500 bp upstream region of the transcription start site as was previously done by Aibar, *et al.* [64], which considers core and proximal promoters. The dot product of the two aforementioned matrices results in a gene-by-TF matrix. We then construct a hypergraph by treating the gene-by-TF matrix as the incidence matrix, where each column that corresponds to a TF is treated as a hyperedge, and genes with elements of “1” in this column are nodes connected by this hyperedge.

The expression matrix of Slide-seq is equivalent to that of scRNA-seq data. Therefore for each Slide-seq cell cluster, we construct an unweighted undirected co-expression k-NN graph using the same method used for scRNA-seq data. In addition, we construct another spatial co-expression graph for each bead cluster which links co-expressed genes in neighboring beads. For this, we identify the neighbors of each bead using the Delaunay triangulation of the bead’s spatial coordinates. Then for each cluster, we compute the correlation of each pair of genes, where one side is the expression of a gene in this cluster and another side is the mean expression of a gene in the neighboring clusters. Again, each gene is connected to the top 10 most co-expressed genes and the graph is converted to a hypergraph where each neighboring-cell gene is treated as a hyperedge.

Finally, we constructed 701 graphs with 1,318,103 nodes in total for 10 tissues and 3 data modalities.

### Connecting gene graphs through a cell cluster dendrogram

We build a cell cluster dendrogram using hierarchical clustering [65], where cell clusters that are close together in the dendrogram are likely functionally similar. For this, we first construct a distance matrix between cell clusters using the obtained top 50 differentially expressed or active genes of each cell cluster. Each Slide-seq cell cluster has two copies, one for the expression graph and one for the spatial graph. Given two sets of differentially expressed or active genes *S_A_* and *S_B_* of cell clusters *A* and cell cluster *B*, their distance is defined as:

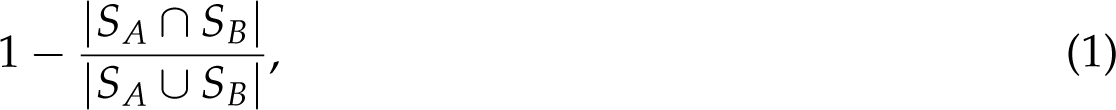

where 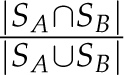 is the Jaccard index of the two gene sets. That is, if two cell clusters have totally different differentially expressed or active gene sets, their distance will be 1, while if they have the same gene sets, their distance will be 0. We then build the dendrogram from the distance matrix using the agglomerative clustering algorithm with the criterion of minimizing the average of distances when merging pairs of branches. The clustering process results in a dendrogram where each leaf is associated with a gene graph of a cell cluster.

### Learning gene embeddings from gene graphs and the cell cluster dendrogram

To identify genes with similar functions or genes participating in the same pathway, we aim to embed each gene in each gene graph into a *d*-dimensional feature space in an unsupervised fashion. For this, we use the multi-layer graph learning objective proposed by Zitnik, *et al.* [22], which relies on connections between genes within a single graph and combines this with inter-graph relationships between genes in different graphs. Formally, let *V* be a given set of *N* genes *{v*_1_, *v*_2_, …, *v_N_}*, and *G_i_* = (*V_i_*, *E_i_*) be one of the *K* gene graphs in the dataset (*i* = 1, 2, …, *K*), where *E_i_*denotes the edges between nodes *V_i_ ⊆ V* within the graph. The dendrogram is a full binary tree *T* defined over a set of 2*K −* 1 elements, where each of the *K* leaf elements is a gene graph. We define *π*(*i*) as the parent of the element *i* in the tree, and reversely, *c*(*i*) as the children of the element *i* in the tree. The problem is thus to learn mapping functions *f*_1_, *f*_2_, …, *f_K_*, each attached with one graph, such that function *f_i_* maps nodes in *V_i_* to feature representations in R*^d^*. *f_i_* is equivalent to a matrix of *|V_i_| × d* parameters.

We started from the assumption that genes with similar neighborhoods in each single graph perform similar functions and therefore should be embedded closely together. The following objective function is used to achieve this goal:

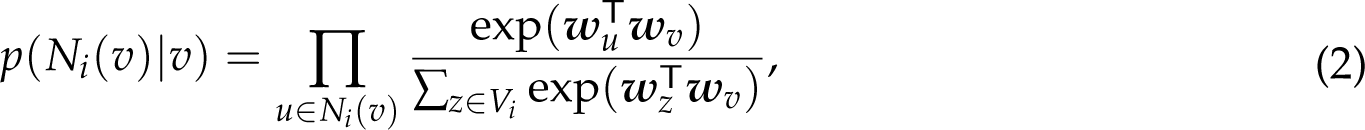

where *N_i_*(*v*) are the neighbors of node *v* in graph *G_i_*, and ***w****_v_* = *f_i_*(*v*), which is the embedding of node *v* in this graph. The conditional likelihood *p*(*N_i_*(*v*)*|v*) is equivalent to the product of a series of softmax functions, maximization of *p*(*N_i_*(*v*)*|v*) therefore tries to maximize the classification probabilities of *v*’s neighbors based on learned node embeddings. With that, the objective for each graph *i* is set to:

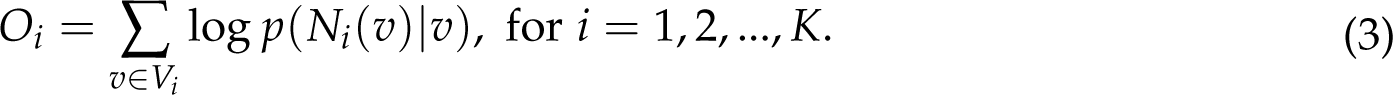

We assume that nearby cell clusters in the dendrogram are functionally similar, which implies that the same gene should perform similar functions in these cell clusters and therefore its embeddings in the corresponding graphs should be similar. For this, we introduce a mapping function *f_i_* (where *i* = *K* + 1, …, 2*K −* 1) for each internal, *i.e.*, non-leaf, element, similar to those of leaf elements, which maps nodes to feature representations in R*^d^*. Recursively, the node set in an internal element is the union of the nodes in its children elements, *V_i_* = *∪_j∈c_*_(_*_i_*_)_*V_j_* (where *i* = *K* + 1, …, 2*K −* 1). The following regularization term is then introduced for nodes in each element *i* of *T*:

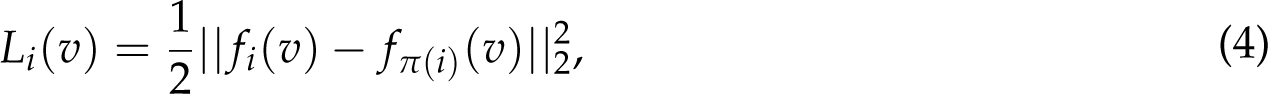

where node *v* in the child elements is encouraged to share similar embeddings with the node *v* in the parent element under the Euclidean norm. We obtain the regularization for an element by summing over the terms of all its nodes: *L_i_*= ∑*_v∈Vi_ L_i_*(*v*). For an internal element *i*, its mapping function *f_i_* can be solved by the following closed-form solution given the mapping functions of all the other elements in *T*, which minimizes the regularization terms involving *f_i_*:

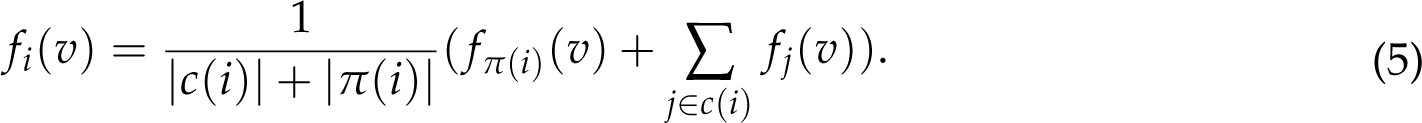

With the objective function and regularization term defined, the model tries to solve the following optimization problem to find the best mapping functions *f*_1_, *f*_2_, …, *f_K_* of leaf elements:

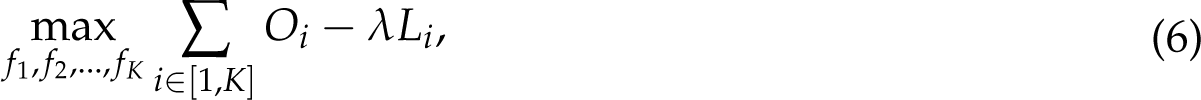

where *λ* is the hyperparameter of the regularization strength. To optimize this function, an iterative learning process is performed, where leaf mapping functions and internal mapping functions are updated alternately. The leaf mapping functions *f*_1_, *f*_2_, …, *f_K_* are updated by performing one epoch of stochastic gradient descent that optimizes Equation (6), while the internal mapping functions *f_K_*_+1_, *f_K_*_+2_, …, *f*_2*K*_*_−_*_1_ are updated using Equation (5).

In practice, we set the dimension hyperparameter *d* as 128 and the regularization strength *λ* as 1.0, which make the model achieve the best performance. When using a *λ* in the range between 1.0 to 10.0, the model performs better by merging information between gene graphs instead of focusing on single gene graphs (*λ* in the range of 0.0 to 0.1). In addition, we observed that the results stay about the same across a range of choices of hyperparameter values for *d* and *λ* (Supplementary Fig. 7). The neighborhood *N*(*v*) for each node in each graph is constructed using Node2vec’s random walk algorithm [66]. We implemented distributed training which applies gradients on each node parallelly based on the Gensim package [67] written in Cython. With this, the algorithm is scalable to hundreds of graphs and millions of nodes.

### Splitting embedding space into components and identifying enriched GOs and TF regulons in them

The learned mapping functions map each gene in each cell cluster to a point in the embedding space. To identify overlapping data sources and discover functions and pathways enriched in different regions of the space, we split the embedding space into embedding components by clustering gene points in it. To do the clustering, we generate another graph in which each point in the space (gene) is connected to its 30 nearest neighbors. The Leiden algorithm [30] with the resolutions of 1 (for identifying overlapping data sources) or 50 (for discovering enriched functions) is then applied to cluster the genes. GO enrichment is performed for genes in each component against all the genes clustered using the GOATOOLS package [68], in which a gene from a cell cluster is considered as an object, and all the genes from all the cell clusters form the population. *P*-values from randomization experiments follow a uniform distribution, indicating the validity of this statistical test (Supplementary Figure 8). GO annotations with the evidence code “IEA”, *i.e.*, electronic annotation without manual review, are ignored in the analysis to avoid circular reasoning. GO annotations are propagated through the GO hierarchies with “is a” and “part of” relationships. False discovery rate (FDR) correction with Benjamini/Hochberg is applied to adjust the enrichment *P*-values.

We also download manually curated TF-target regulatory relationships from the TRUST v2 [69] database. Enrichment of TF regulons (genes regulated by the same transcription factor) in each component is computed using the same method described above.

### Inferring multicellular functions of genes

To identify new functions for genes we relied on genes with known functions that are co-embedded with them. Specifically, for one gene, we calculate the Euclidean distance between each pair of the gene’s embeddings of different graphs, and then group all its graphs into clusters using agglomerative clustering, where two clusters will stop being merged if the distance between them is above 1.5. Then for each graph in a cluster, the gene’s 50 nearest neighbors in the graph are identified based on the Euclidean distance between gene embeddings. A “common neighbor” set for the gene in this cluster is finally selected based on two criteria: (1) the neighbor gene must appear in the nearest neighbors of at least half of the graphs in the cluster; (2) the neighbor gene is in the 50 most frequent neighbors based on the number of its appearance in nearest neighbors. Next, GO enrichment analysis is performed using the common neighbors. GO terms that are enriched in the neighbors (*P <* 0.05) are assigned to the gene as predicted functions. In this analysis, we only predict functions of genes expressed in at least 30% of the cells in each cell cluster.

### Complete GIANT workflow

Below is the description of the full GIANT workflow, which is implemented as the Python interface:

1. Read in gene expression matrices (scRNA-seq and Slide-seq) and gene activity matrices (scATAC-seq) of all the tissues. Identify the union of variable genes of each tissue.
2. Read in scRNA-seq gene expression matrix of each tissue. Construct a co-expression graph for each cell cluster on the variable genes.
3. Read in peaks for each cell cluster of each tissue, genomic regions containing TF motifs, and gene coordinates. Construct a gene-TF hypergraph for each cell cluster on the variable genes.
4. Read in Slide-seq gene expression matrix of each tissue, and locations of beads. Construct a co-expression graph and a spatial co-expression graph for each cell cluster on the variable genes.
5. Read in differential genes of each cell cluster and construct a cell cluster dendrogram.
6. Run gene embedding.
7. Cluster genes in the entire embedding space into clusters (embedding components).
8. Run GO enrichment in each embedding component.
9. Run TF regulon enrichment in each embedding component.
10. Cluster embeddings of each gene into clusters, and identify the gene’s neighbors in each cluster.
11. Inferring functions of each gene in each cluster by running GO enrichment in the neighbor genes.

### Data availability

The gene embeddings generated from GIANT for both the HuBMAP dataset and the human fetal atlas dataset are available at https://github.com/chenhcs/GIANT. Accession numbers for the HuBMAP datasets used in this study are listed in Supplementary Table 4. Data can be obtained from the HuBMAP data portal (https://portal.hubmapconsortium.org) via accession numbers. The scRNA-seq data and the scATAC-seq data of the human fetal atlas were obtained from the descartes interactive website (https://descartes.brotmanbaty.org/).

### Code availability

The source code of GIANT is available at https://github.com/chenhcs/GIANT.

## Acknowledgements

The work in this article was partially supported by National Institutes of Health (NIH) grants OT2OD026682, 1U54AG075931 and 1U24CA268108 to Z.B.-J.

## Author contributions

H.C., N.D.N. and Z.B.-J. conceptualized the study. H.C., N.D.N. and Z.B.-J. designed the algorithm and methodology. M.R. and H.C. performed data preprocessing. H.C. and N.D.N. developed the software. H.C. and Z.B.-J. performed the analysis. H.C., N.D.N. and Z.B.-J. wrote the manuscript. All authors read and approved the final manuscript.

