## Supplementary Notes and Figures for "A unified analysis of atlas single cell data"

#### Silhouette coefficient for data modalities

We calculated the mean silhouette coefficient over all the nodes in the t-SNE space of each embedding by considering their source of data modalities. Specifically, the silhouette Coefficient was first calculated for each node:

$$\frac{b-a}{\max(a,b)},$$

where $a$ is the mean intra-modality distance for the node and $b$ is the mean nearest-modality distance for the node, defined as the distance between a node and the nearest modality that the node is not a part of. The mean Silhouette Coefficient over all nodes is then calculated, which ranges between -1 and 1. A silhouette coefficient near 0 indicates different data modalities are overlapping globally, while a silhouette coefficient near 1 indicates nodes are clustered based on their modalities. We consider Slide-seq expression and Slide-seq spatial as one modality to avoid considering the trivial case where two representations of the same data overlap. Thus there are three groups, scRNA-seq, scATAC-seq, and Slide-seq, in the calculation of silhouette coefficients.

#### Comparisons with cell-based embedding methods

To compare with GIANT, we performed cell embeddings on the HuBMAP dataset using five popular cell-based embedding methods, including Harmony [1] (clustering based correction), LIGER [2] (non-negative matrix factorization based), Scanorama [3] (mutual nearest neighbor based), scVI [4] (deep learning based), Seurat v4 [5] (mutual nearest neighbors based), and GLUE (deep learning based) [6]. We note that there are many other methods developed for multi-modality single cell data integration, however, may not be suitable to be applied to the datasets we used. For example, both multiVI [7] and MOFA+ [8] require input data where multi-omics measurements are derived from the same set of cells. Such data are not available in our dataset.

For Harmony, LIGER, Scanorama, scVI, and Seurat, the gene count matrix for scRNA-seq and gene activity matrix for scATAC-seq were used as input as suggested by Seurat and LIGER [2, 5]. For GLUE, the gene count matrix for scRNA-seq and the cell-by-bin count matrix for scATAC-seq were used as input, and the default regulatory graph was constructed to connect ATAC bins to genes if they overlap in either the gene bodies or promoter regions. The reciprocal PCA method was used for Seurat to improve speed. The parameters of different methods were set as default. We computed the same silhouette coefficient for each of the cell-based methods.

To evaluate the ability of the cell-based method in discovering functional gene modules across datasets, for each cell-based method, we clustered cells into cell clusters based on its embeddings using the Leiden algorithm [9] with the resolution of 10. Then a co-expression network is built for each cell cluster using the same method as in GIANT. The WGCNA module detection method [10] is then applied to detect gene modules from each co-expression network by hierarchical clustering of genes and dynamic tree cut. GO enrichment was then performed in each gene module. We aggregated the enriched GO terms for each method.

The GLUE method also generates gene embeddings along with the cell embeddings, however, there is only one embedding for each gene in the whole dataset (vs. one embedding for each gene in each cell cluster for GIANT), which cannot reflect tissue or cell type-specific functions of the gene. We therefore only compared with the cell embeddings of GLUE.

##### Application on the human fetal atlases

We applied GIANT on another dataset of human fetal atlas that contains scRNA-seq data of 4,062,965 cells from 15 tissues [11], including Adrenal, Cerebellum, Cerebrum, Eye, Heart, Intestine, Kidney, Liver, Lung, Muscle, Pancreas, Placenta, Spleen, Stomach, Thymus, and scATAC-seq data [12] of 720,613 cells from the same 15 tissues.

We used the cell clusters provided by the atlas with cell type annotations. Specifically, cells in the scRNA-seq data were clustered using the Louvain clustering [13] on the UMAP space per tissue. Clusters were annotated based on cell type-specific marker gene expression. Cells in the scATAC-seq data were annotated with cell types by leveraging the annotations on the scRNA-seq data of the same tissues, on the basis of gene-level accessibility scores computed for the scATAC-seq data. We finally got 172 cell clusters of 15 tissues for the scRNA-seq data and 100 cell clusters of 15 tissues for the scATAC-seq data.

The same algorithms were used to construct a gene co-expression graph for each scRNA-seq cell cluster and a gene-TF hypergraph for each scATAC-seq cell cluster. To get the list of peaks for each scATAC-seq cell cluster, specificity scores for peak and cell type pairs were obtained from the original publication [12]. We consider the top 500,000 peaks for each cell type with the highest specificity scores.

To construct the dendrogram, the top 50 differentially expressed or active genes were computed from the gene expression matrix of each scRNA-seq cell cluster and the gene-level accessibility matrix of each scATAC-seq cell cluster, which was computed in the original publication [12].

To make results comparable between the HuBMAP dataset and the human fetal dataset, we consider the same set of genes selected from the HuBMAP dataset for generating the gene embeddings of the human fetal dataset.

After the gene embeddings were obtained. Embedding components were again identified from this human fetal dataset. To assess the consistency between the embedding components obtained from the two datasets, we computed the enrichment of the 1,199 HuBMAP embedding components in the human fetal dataset. Specifically, for each HuBMAP embedding component, we treated the set of genes appearing in the embedding component as a gene module. The enrichment of the gene modules in the set of genes of each embedding component of the human fetal dataset was then computed.

### Supplementary Figures

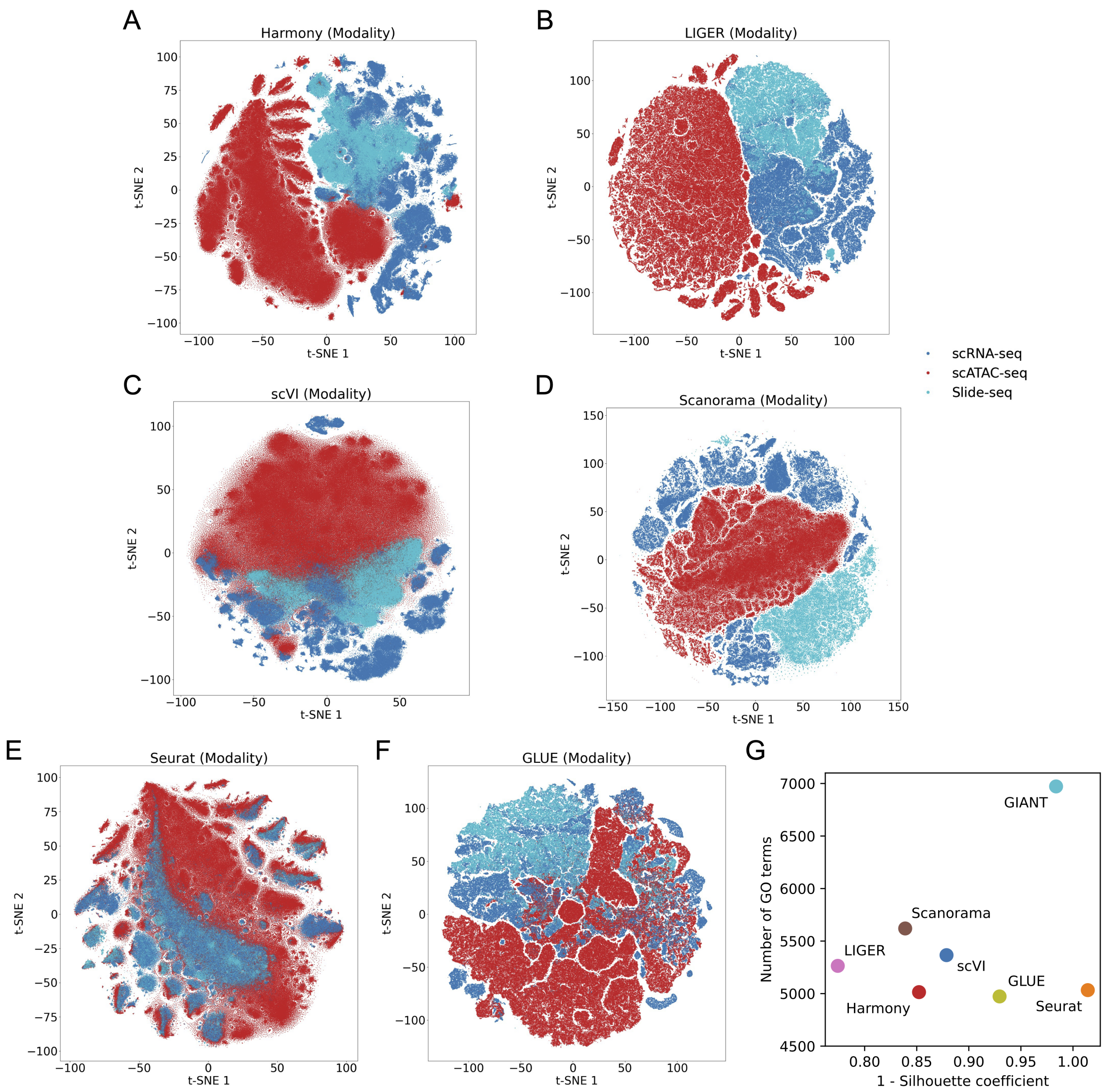


Supplementary Figure 1: (A-F) Visualization of cell embeddings on the HuBMAP dataset from six cell-based methods. Each point in the visualization represents a cell. Cells are colored by their data modalities. (G) The comparisons between the GIANT gene embeddings and the other cell-based embeddings. More unique GO terms are found enriched in the GIANT embedding components than in the gene modules identified from the competing cell-based embeddings. For GIANT, GO enrichment because of multiple copies of a single gene is ignored. On the other hand, GIANT shows a larger 1 - silhouette coefficient for data modalities than most of the cell-based embeddings, indicating more overlapping between different modalities globally.

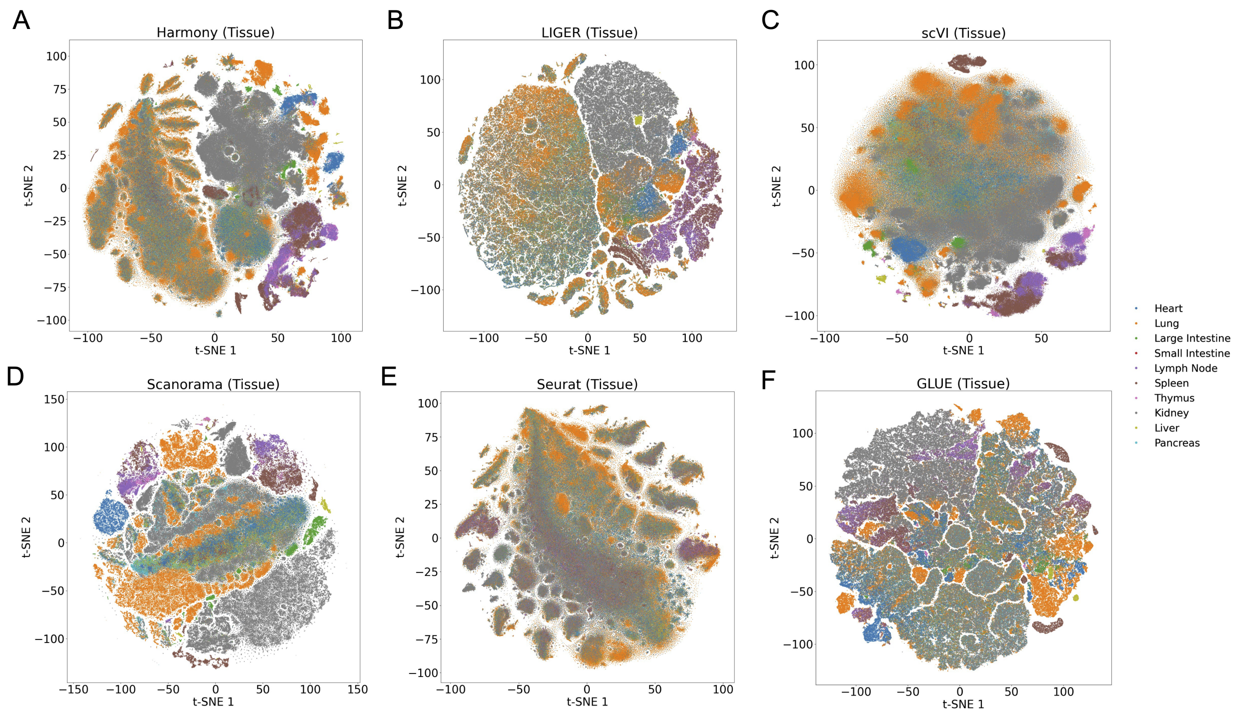


Supplementary Figure 2: (A-F) The visualization of cell embeddings on the HuBMAP dataset for six cell-based methods (indicated in the titles of subfigures). Each point in the visualization represents a cell. Cells are colored by their tissues.

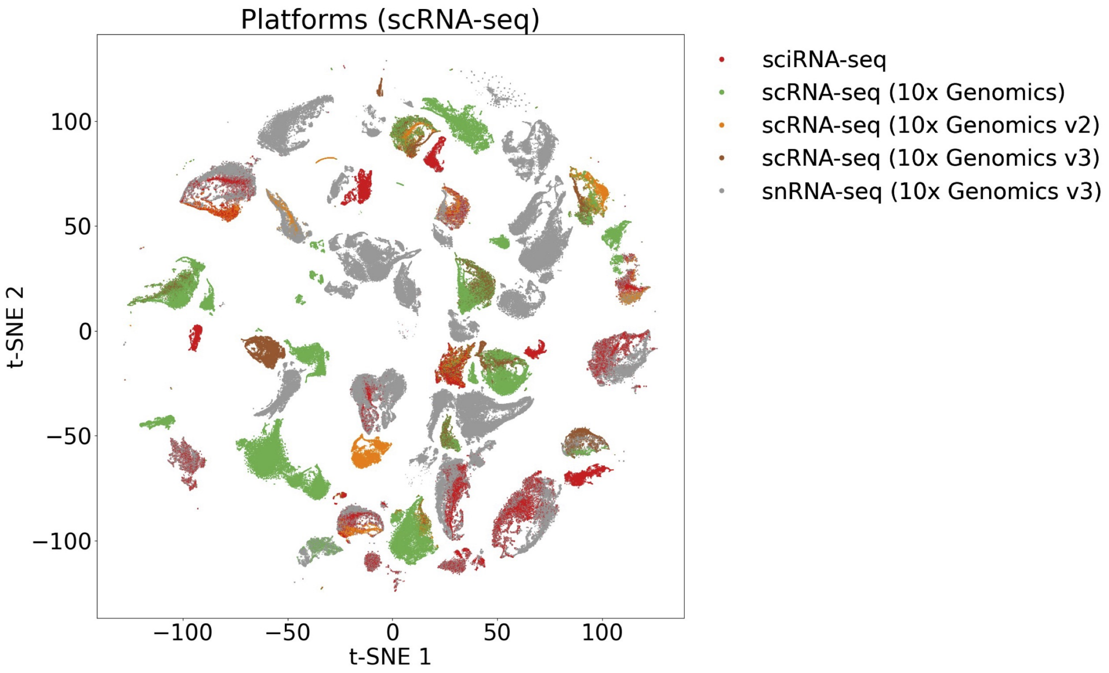


Supplementary Figure 3: Visualization of GIANT gene embeddings on the HuBMAP dataset. Only the scRNA-seq modality is shown in this plot. Genes are colored by the platforms that the datasets were generated from.

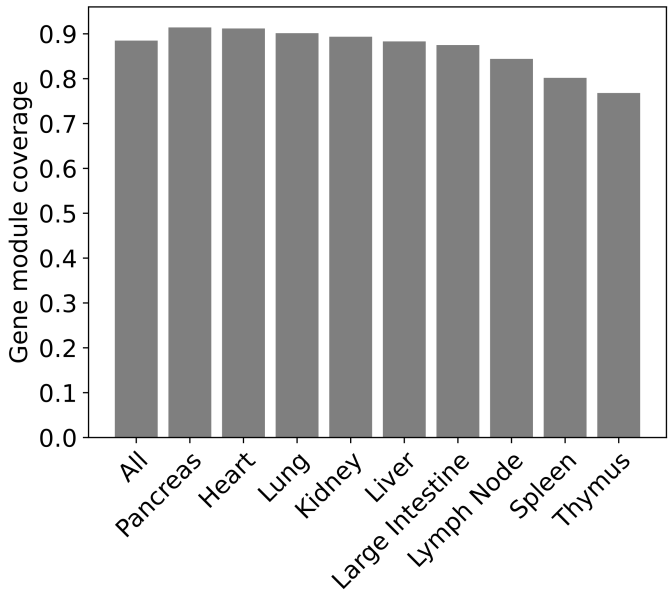


Supplementary Figure 4: The percentage of gene modules associated with each HuBMAP human tissue that have enrichment in the embedding components of the human fetal atlas dataset.

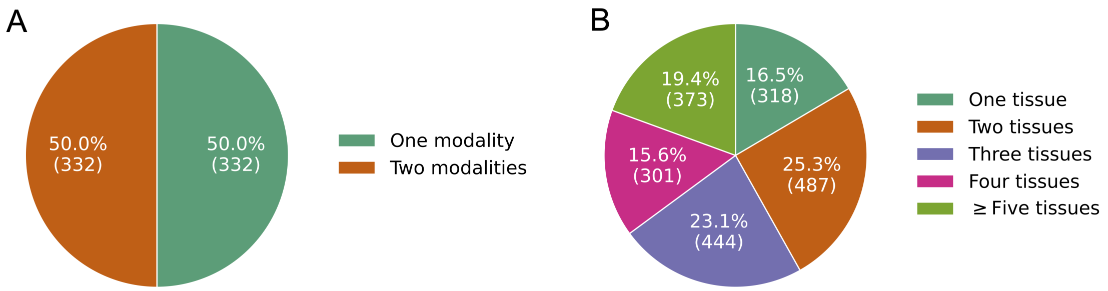


Supplementary Figure 5: (A) The number of genes that have predicted functions arising out of neighbors from one modality or at least two modalities. 664 genes that appear in at least two of the three data modalities are considered. (B) The number of genes that have predicted functions arising out of neighbors from different numbers of tissues. 1,924 genes that appear in at least two tissues are considered.

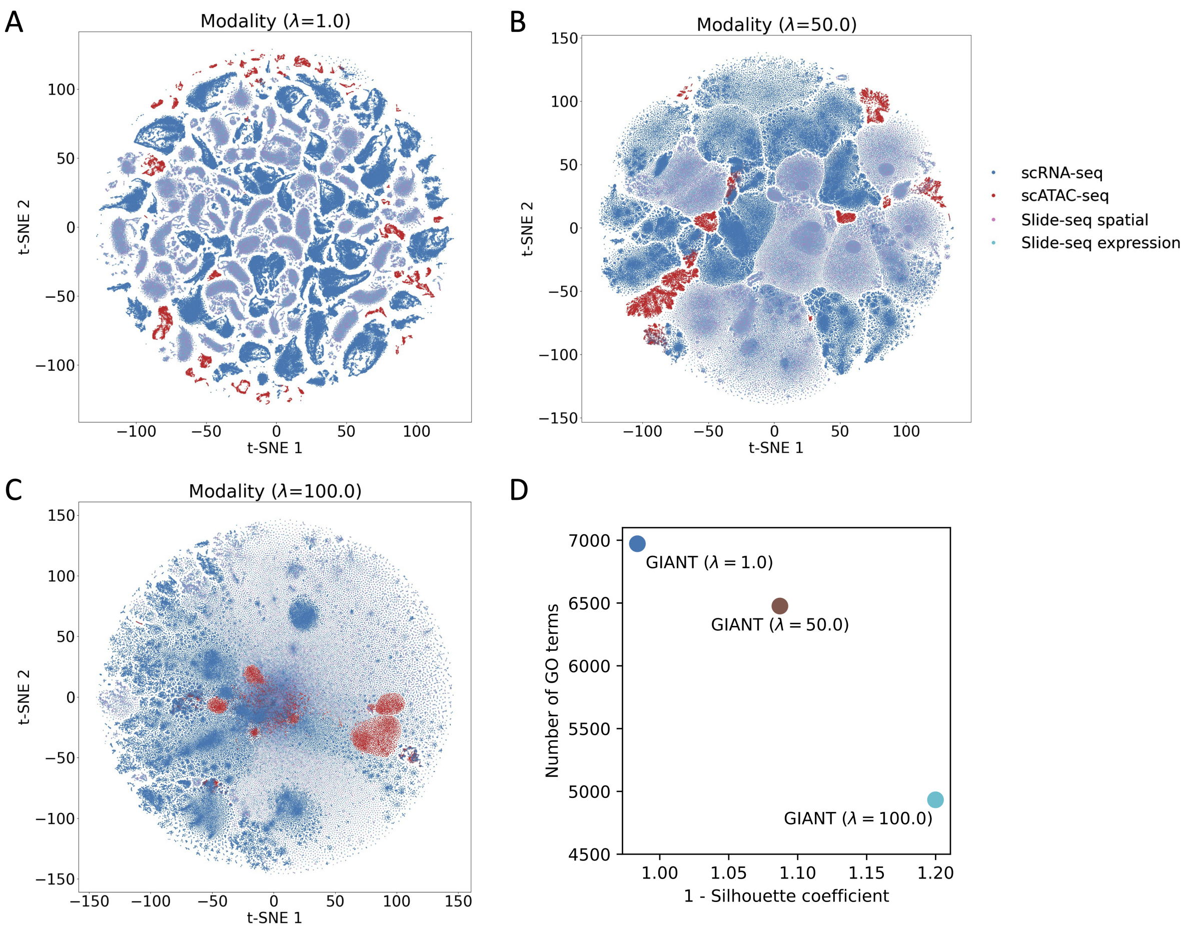


Supplementary Figure 6: (A-C) Visualization of GIANT gene embeddings on the HuBMAP dataset for different regulation strength parameters ($\lambda$). Genes are colored by their data modalities. (D) Larger $\lambda$ can increase the level of modality merging, indicated by the larger 1-Silhouette coefficients, however, the GO terms that are found enriched in the embedding components are decreased, which indicates lower qualities of these embedding components.

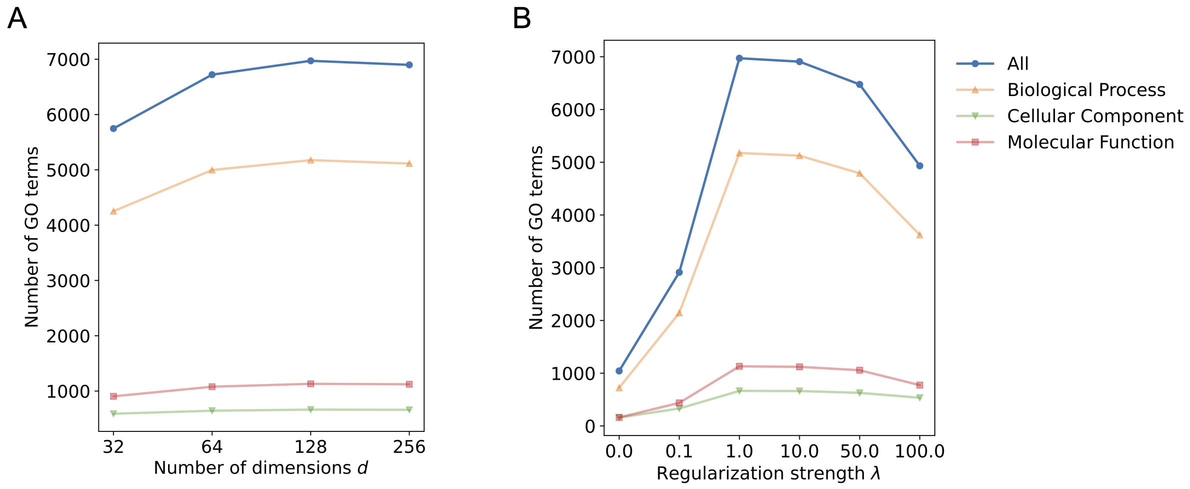


Supplementary Figure 7: Effect on the performance of GIANT of the choices of hyperparameters for the embedding dimensions ($d$) and regulation strength for balancing two learning objectives ($\lambda$). We measure the performance using the number of different types of GO terms that can be discovered from different embedding components (Methods). (A) While the performance improves when the number of dimensions is up to 128, the performance is overall stable across a reasonable range of values. (B) The performance drops when the model focuses on only the single graph learning objective ($\lambda = 0.0$) or applies too much regularization, though when $\lambda$ is around 1.0 to 10.0, the model achieves the best performance.

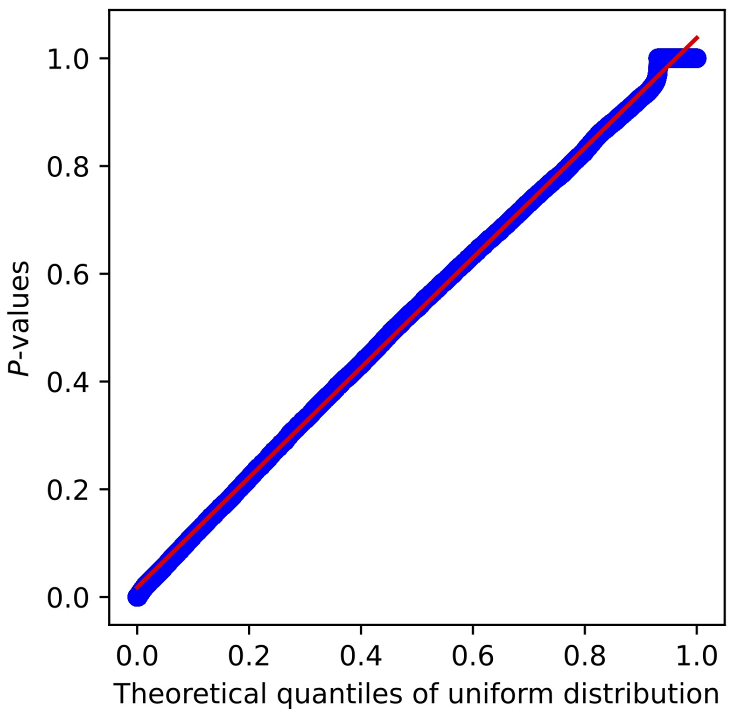


Supplementary Figure 8: The quantiles of the distribution of $P$-values against those of uniform distributions. The plot indicates the distribution of $P$-values is uniform.
